# Nuclear genome of a pedinophyte pinpoints genomic innovation and streamlining in the green algae

**DOI:** 10.1101/2021.10.04.463119

**Authors:** Sonja I Repetti, Cintia Iha, Kavitha Uthanumallian, Christopher J Jackson, Yibi Chen, Cheong Xin Chan, Heroen Verbruggen

## Abstract

The genomic diversity underpinning high ecological and species diversity in the green algae (Chlorophyta) remains little known. Here, we aimed to track genome evolution in the Chlorophyta, focusing on loss and gain of homologous genes, and lineage-specific innovations of the Core Chlorophyta. We generated a high-quality nuclear genome for pedinophyte YPF701, a sister lineage to others in the Core Chlorophyta, and incorporated this genome in a comparative analysis with 25 other genomes from diverse Viridiplantae taxa. The nuclear genome of pedinophyte YPF701 has an intermediate size and gene number between those of most early-diverging prasinophytes and the remainder of the Core Chlorophyta. Our results suggest positive selection for genome streamlining in Pedinophyceae, independent from genome minimisation observed among prasinophyte lineages. Genome expansion was predicted along the branch leading to the UTC clade (classes Ulvophyceae, Trebouxiophyceae and Chlorophyceae) after divergence from their common ancestor with pedinophytes, with genomic novelty implicated in a range of basic biological functions. These results emphasise multiple independent signals of genome minimisation within the Chlorophyta, as well as the genomic novelty arising prior to diversification in the UTC clade, which may underpin the success of this species-rich clade in a diversity of habitats.

## Introduction

The Chlorophyta are a diverse group of green algae, belonging, along with the Streptophyta and Prasinodermophyta, to the Viridiplantae, an ancient lineage that diverged from a putative ‘ancestral green flagellate’ (Leliaert *et al*., 2012; Fang *et al*., 2017; Li *et al*., 2020). The Chlorophyta are subdivided into the Core Chlorophyta and the paraphyletic early branching prasinophytes, which are mostly marine unicellular planktonic algae (Marin, 2012; Fučíková *et al*., 2014; Fang *et al*., 2017, 2018). The more species-rich Core Chlorophyta comprise the well-supported ‘UTC’ clade - which is composed of the classes Ulvophyceae, Trebouxiophyceae and Chlorophyceae - and the smaller and earlier diverging Chlorodendrophyceae and Pedinophyceae (Del Cortona *et al*., 2020).

Numerous studies (e.g. Derelle *et al*., 2006; Palenik *et al*., 2007) have analysed both the coding and noncoding elements in the nuclear genomes of prasinophyte lineages, elucidating features correlated with early divergence and diversification of green algae (Lemieux *et al*., 2019). The prasinophyte genus *Ostreococcus* represents some of the smallest free-living eukaryotes with relatively small (∼13 Mb) genomes (Derelle *et al*., 2006; Palenik *et al*., 2007). Compared to genomes of other chlorophytes, the reduced genomes of prasinophytes exhibit smaller numbers of gene families and genes, shortened intergenic regions, and fused genes (Derelle *et al*., 2006; Moreau *et al*., 2012). The small and gene-dense genomes of prasinophytes may reflect genome streamlining, a hypothesis that postulates selection acts to minimize the cost of replicating non-essential DNA, thereby reducing genome size (Giovannoni, 2005). Studies have concluded that genome minimization has occurred separately in prasinophyte groups Chloropicophyceae and Mamiellophyceae, involving different predicted losses of genes and pathways (Lemieux *et al*., 2019).

Most genomic studies in the Core Chlorophyta have investigated taxon-specific innovations that underpin their ecological success under a range of environmental pressures including high acidity (Hirooka *et al*., 2017), high salinity (Foflonker *et al*., 2015), polar conditions (Blanc *et al*., 2012; Zhang *et al*., 2020), and as symbionts (Blanc *et al*., 2010; Arriola *et al*., 2018; Iha *et al*., 2021). Members of the Core Chlorophyta, particularly the UTC clade, also show high morphological diversity (Table S1), including unicellular, siphonous, and multicellular forms, which appear to have arisen on multiple occasions in both the Chlorophyceae and Ulvophyceae (Featherston *et al*., 2017; De Clerck *et al*., 2018; Del Cortona *et al*., 2020). Although the number of sequenced genomes for the Core Chlorophyta is increasing steadily, the genomic diversity underpinning their ecological and species diversity remains to be systematically investigated.

Positioned as sister to the rest of the Core Chlorophyta (Del Cortona *et al*., 2020), the class Pedinophyceae (pedinophytes) (Moestrup, 1991; Marin, 2012) presents an excellent study subject to examine the evolution of the Core Chlorophyta, including the gene family evolution that occurred as this group diverged. Pedinophytes are small (2.5-7.0 μm), usually naked, unicellular green flagellates found in water or soil habitats and sometimes in symbioses (Sweeney, 1976; Cachon & Caram, 1979; Karpov & Tanichev, 1992; Marin, 2012; Jackson *et al*., 2018). Pedinophyte morphology varies greatly, and they have been described in a variety of environments ranging from freshwater, marine, to hyperhaline (Karpov & Tanichev, 1992; Jones *et al*., 1994).

In this study, we present a high-quality nuclear genome of a pedinophyte. Incorporating this genome in a comparative analysis with 25 genomes from other Viridiplantae taxa, we investigated genome evolution in the Chlorophyta, focussing on patterns of gene-family loss and gain. We emphasise the implication of these results for innovations that may have emerged upon divergence and subsequent diversification of the Core Chlorophyta lineage.

## Materials and Methods

### Culturing and nucleic acid extraction

Pedinophyte strain YPF701 (NIES Microbial Culture Collection strain NIES-2566) was cultured in K-enriched seawater medium (Keller *et al*., 1987) at 20 °C on a 10:14 hour light:dark cycle. To reduce bacterial load, cultures were treated with antibiotics (cefotaxime 0.72mg/mL, carbenicillin 0.72mg/mL, kanamycin 0.03mg/mL and amoxicillin 0.03mg/mL) one week prior to extraction for long-read sequencing. Total genomic DNA was extracted using a modified CTAB protocol, in which the CTAB extraction buffer was added directly to the cell pellets (Cremen *et al*., 2016).

### DNA and RNA Sequencing

DNA was sequenced using nanopore sequencing technology (MinION, Oxford Nanopore Technologies), producing approximately 953,000 reads and 6.61 GB, with an average read length of approximately 7,000 bp.

DNA was prepared for short-read sequencing using a Kapa Biosystems kit, for sequencing of 2 × 150 bp paired-end reads using the Illumina NextSeq platform at Novogene, Hong Kong (see Jackson *et al*., 2018). For RNA sequencing, total RNA was extracted using PureLink™ Plant RNA Reagent (Thermofisher, Waltham, MA, USA). A strand-specific 100 bp paired-end library was constructed and sequenced using Illumina HiSeq 2500.

### *De novo* assembly of pedinophyte transcriptome and genome

Removal of adaptors from long-read data was performed with Porechop (Wick *et al*., 2017), https://github.com/rrwick/Porechop). Quality filtering was performed using Filtlong (https://github.com/rrwick/Filtlong), with quality cutoff 50 and window quality cutoff 40.

Quality filtering of short reads was performed using Trimmomatic v0.39 (Bolger *et al*., 2014) with the following settings: LEADING:3 TRAILING:3 SLIDINGWINDOW:4:20.

Transcriptome data were assembled using Trinity v2.8.3 (Grabherr *et al*., 2011).

*De novo* genome assembly was conducted using MaSuRCa v3.2.8 (Zimin *et al*., 2013) at default setting, combining both long- and short-read data.

The assembled genome was filtered to remove contaminants based on a comprehensive strategy employing taxonomic annotations, read coverage, transcriptome data, and GC content. For this process, genes were predicted for the scaffolds using GeneMark-ES version 2.0 (Ter-Hovhannisyan *et al*., 2008), and the associated coding sequences (CDS) were searched (BLASTN) against the GenBank nucleotide (nt) database and subsequently categorized as green algae, bacteria, or other, based on the top hit. We manually verified this categorisation and removed scaffolds with a high similarity to sequenced bacterial genomic data, no predicted genes, no mapped transcripts, and/or deviant average coverage of sequencing reads or GC content from the assembly. This ensured high confidence that retained scaffolds represent correctly assembled segments of the pedinophyte genome. Scaffolds corresponding to the mitochondrial and chloroplast genomes were also removed based on their CDS matching sequenced organelle genomes on GenBank. This approach revised the assembled genome from 34 Mbp (1877 scaffolds) to the final assembly of 28 Mbp (32 scaffolds).

Genome summary statistics were calculated with QUAST 5.0.2 (Mikheenko *et al*., 2018) and Geneious 11.1.2 (Kearse *et al*., 2012).

### *Ab initio* prediction of protein-coding genes

After filtering of scaffolds, we followed the workflow described in Iha *et al*. (2021) to predict protein-coding genes from the assembled genome sequences. Novel repeat families were identified with RepeatModeler v1.0.11 (http://www.repeatmasker.org/RepeatModeler/). All repeats (including known repeats in RepeatMasker database release 20181026) in the genome scaffolds were masked using RepeatMasker v4.0.7 (http://www.repeatmasker.org/) before gene prediction.

To generate high-quality evidence to guide gene prediction, we first employed PASA pipeline v2.3.3 and TransDecoder (Haas *et al*., 2003) to predict transcript-based protein-coding genes from the unmasked genome assembly and the assembled transcriptome. Predicted proteins were searched (BLASTP, *E* ≤ 10^−20^, >80% query cover) against proteins in RefSeq database (release 88), and checked for transposable elements using HHblits v2.0.16 (Remmert *et al*., 2012) and TransposonPSI (Haas, 2007). Predicted proteins with hits to RefSeq and no transposable elements were retained, and redundant sequences were removed using CD-HIT v4.6.8 (Li & Godzik, 2006) (-c 0.75 -n 5). The resulting gene models were then used to infer high-quality “golden genes” using the script *Prepare_golden_genes_for_predictors*.*pl* from the JAMg pipeline (https://github.com/genomecuration/JAMg). These “golden genes” were used as the training set to guide gene prediction in the repeat-masked genome sequences with AUGUSTUS v3.3.1 (Stanke *et al*., 2006) and SNAP (Korf, 2004). Additional gene models were generated with GeneMark-ES v4.38 (Lomsadze *et al*., 2005) and MAKER v2.31.10 (Holt & Yandell, 2011) (protein2genome, UniProt-SwissProt database retrieved 27 June 2018). Protein-coding genes predicted using the five methods (PASA, AUGUSTUS, SNAP, MAKER, and GeneMark-ES) were integrated using EvidenceModeler v1.1.1 (Haas *et al*., 2008). The weights for each gene prediction output were: GeneMark-ES 2, MAKER 8, PASA 10, SNAP 2, AUGUSTUS 6. We retain PASA-predicted genes (which are supported by transcriptome evidence), and those predicted by two or more other methods, as the final set of protein-coding genes.

### Comparison of nuclear genomes for the green lineage

For comparative genomic analyses, we built a dataset, containing both genomes and proteomes, of 20 Chlorophyta taxa, including pedinophyte strain YPF701, 5 Streptophyta taxa, and *Prasinoderma coloniale* (Table S1). Percentage of identified BUSCO sequences was assessed for all proteomes with BUSCO v5.2.2 (Manni *et al*., 2021), using the chlorophyta_odb10 lineage for members of the Chlorophyta, streptophyta_odb10 lineage for members of the Streptophyta and viridiplantae_odb10 for *Prasinoderma coloniale* and *Klebsormidium nitens*.

GC content of CDS and synonymous codon usage order (SCUO) were calculated using the CodonO (Angellotti *et al*., 2007) function from the cubfits v.0.1-3 (Chen *et al*., 2014) package in R version 3.5.1 (R Core Team, 2020). SCUO value ranges from 0 to 1, with a larger value indicating stronger codon usage bias.

We used the OrthoFinder 2.5.1 (Emms & Kelly, 2019) pipeline (default parameters) to cluster proteins from the dataset into homologous groups (i.e. “orthogroups” defined by the program).

A phylogenetic tree was manually constructed to reflect current knowledge of evolutionary relationships between taxa from large-scale multi-gene phylogenies (Fig. 1) (Del Cortona *et al*., 2020; Li *et al*., 2020).

**Fig. 1.**
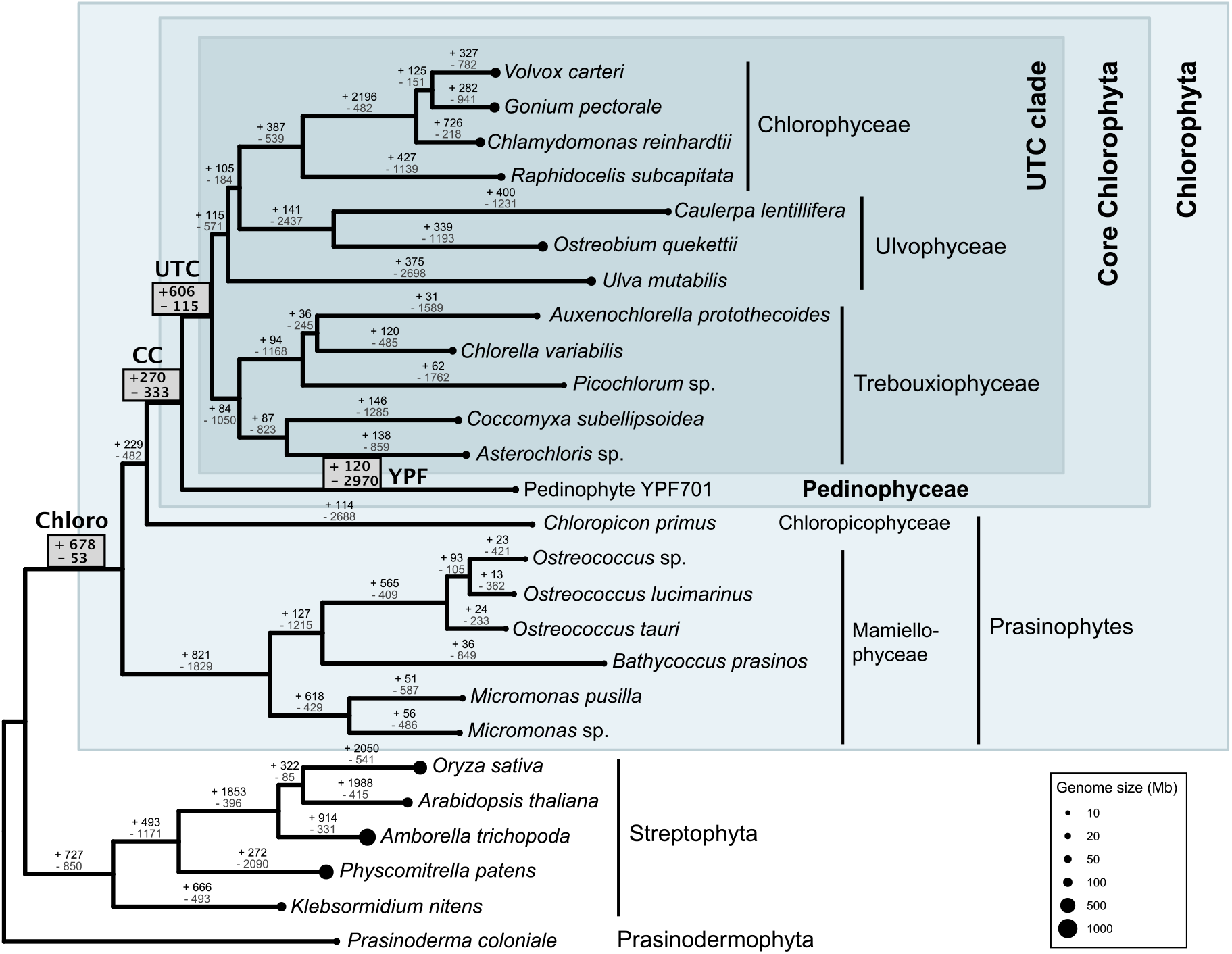
Phylogenetic tree of green algae and land plants, including predicted pattern of gain and loss of orthogroups. The number of gene families acquired (black) or lost (dark grey), indicated along each branch in the tree were estimated using the Dollo parsimony principle. Size of the circle at branch tips is proportional to genome size. Branches of interest in our study are marked in bold: base of the Chlorophyta (Chloro), Core Chlorophyta (CC), UTC clade (UTC) and branch to pedinophyte YPF701 (YPF).

Based on the phylogenetic tree, the most parsimonious gain and loss scenario was reconstructed for each orthogroup using the Dollop program from PHYLIP version 3.695 (Felsenstein, 2005), with the Dollo parsimony method and printing of states at all nodes of the tree. This gain and loss scenario was processed using extract_dollop_output_sequences_v2-fast.pl from OrthoMCL Tools v1.0 (Leonard, 2015), and mapped to the tree in RStudio using R version 4.0.2 (R Core Team, 2020) with the packages phytools 0.7.70 (Revell, 2012), ape 5.4.1 (Paradis & Schliep, 2019), maps 3.3.0 (Brownrigg *et al*., 2018), ggplot2 3.3.2 (Wickham, 2016) and ggtree v2.2.4 (Yu *et al*., 2017).

Orthogroup losses and gains were further analysed by examining their annotated Gene Ontology (GO) terms. Chlorophyta proteomes were analysed using eggNOG-mapper 2.0.1 (Huerta-Cepas *et al*., 2017, 2019), with DIAMOND 0.9.24 (Buchfink *et al*., 2015), default settings, and ‘Viridiplantae’ as taxonomic scope to maximise accurate functional annotations. GO terms associated with sequences found in orthogroups gained or lost along branches of interest were summarised using REVIGO (Supek *et al*., 2011), focusing on the ‘Biological Process’ category. REVIGO results were visualised using CirGO (Kuznetsova *et al*., 2019), weighted according to the number of gained/lost orthogroups associated with each GO term, including full eggNOG-mapper results for branches of interest except for orthogroups lost along YPF, for which only the top 3500 GO terms were included (when sorted by number of gained/lost orthogroups associated with each GO term) due to REVIGO constraints.

GO analyses using *Chlamydomonas reinhardtii*, which has more comprehensive GO annotations relative to most Chlorophyta, were also used to explore functions of orthogroups that contained *C. reinhardtii* sequences. Orthogroups gained or lost along branches of interest were grouped into functional clusters according to ChlamyNET (Romero-Campero *et al*., 2016), which provides a gene co-expression network of *C. reinhardtii* transcriptomes.

## Results

The newly assembled nuclear genome for pedinophyte YPF701 comprises 32 scaffolds with a total length of 27,899,919 bp, scaffold N50 of 1.23 Gb, and 7,940 predicted protein-coding genes (Table S1). The genome has a size, number of proteins, and average number of genes per orthogroup that are intermediate between those of most prasinophytes and the rest of the Core Chlorophyta (Fig. 2). The GC content is 70%, which is higher than most sequenced green algal nuclear genomes but not unseen in the Chlorophyta (Suzuki *et al*., 2018). The genome shows the highest synonymous codon usage order of the Viridiplantae genomes included in this study (Fig. 2).

**Fig. 2.**
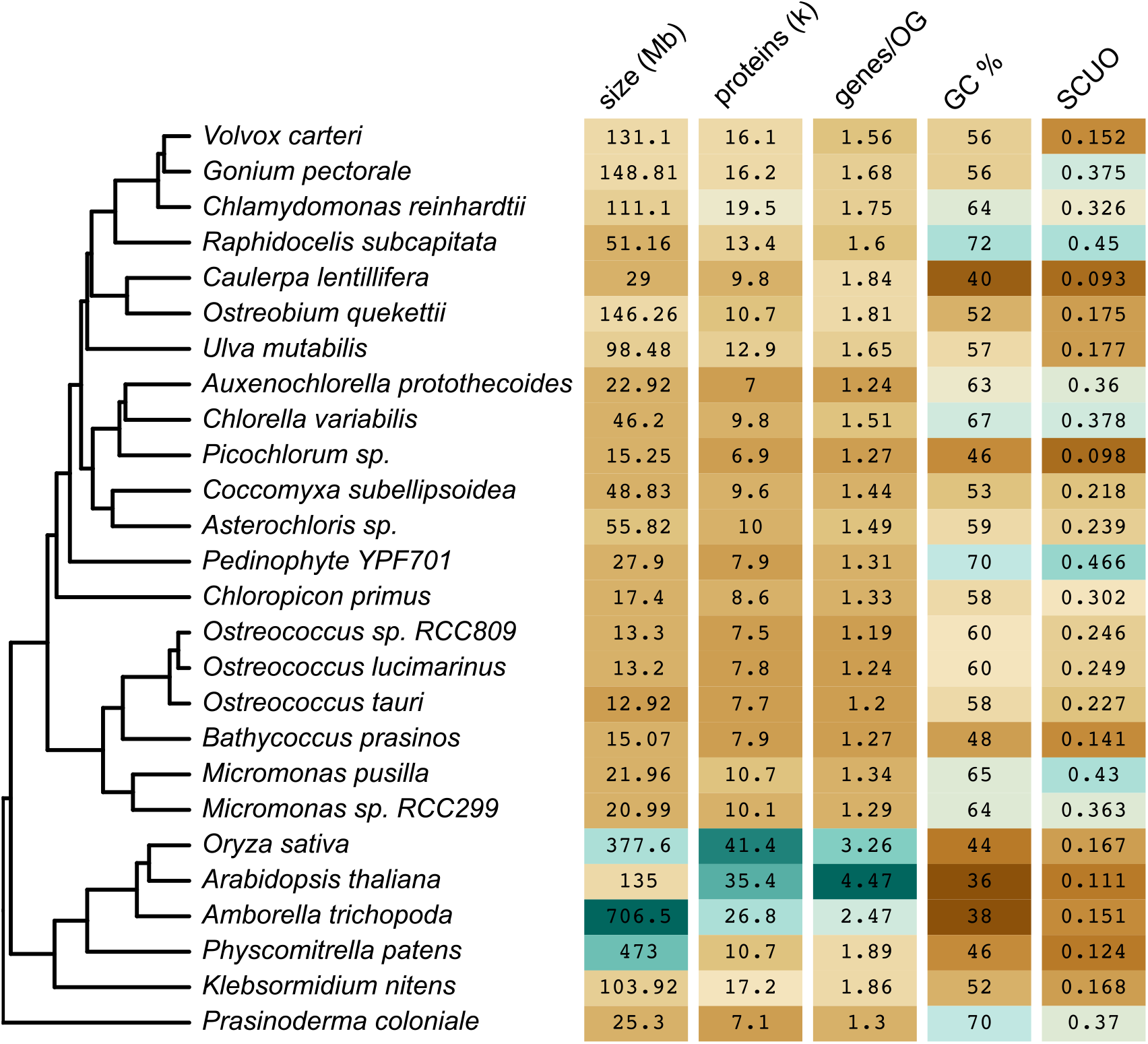
Heat map of key traits for Viridiplantae nuclear genomes used for analysis in this study (brown= lowest value, blue=highest value). SCUO = synonymous codon usage order.

Comparative analysis of predicted proteins from 26 genomes (Table S1) reveals patterns of putative gains and losses of homologous groups (i.e. OrthoFinder-defined “orthogroups”) across the Viridiplantae phylogeny, showing predicted losses outnumbering predicted gains for most Chlorophyta branches (Fig. 1). Associated GO terms and ChlamyNET classifications for sequences suggest potential functions for a subset of the orthogroups gained or lost along the branches at the base of the Chlorophyta (Chloro), Core Chlorophyta (CC) and UTC clade (UTC) and leading to the pedinophyte genome (YPF) (Tables S2, S3), with many GO terms implicated in metabolism and biological processes related to signalling and regulation in cells (Fig. S1). The use of ‘Viridiplantae’ as taxonomic scope for eggNOG-mapper, due to how little GO data is available for the Chlorophyta, resulted in GO term annotations for plant-specific processes, including “pollen tube development” and “regulation of flower development” (Table S2), which likely incorporate these biological functions. Results from ChlamyNet analysis reinforce these functional themes of cell regulation and metabolism (Fig. 3), with the greatest number of ChlamyNet hits for most branches falling into cluster 3: “protein phosphorylation, ribosome biogenesis and macromolecular synthesis”. This is the largest Chlamynet cluster, which is involved in diverse biological processes and is significantly enriched in transcription factors (Romero-Campero *et al*., 2016).

**Fig. 3.**
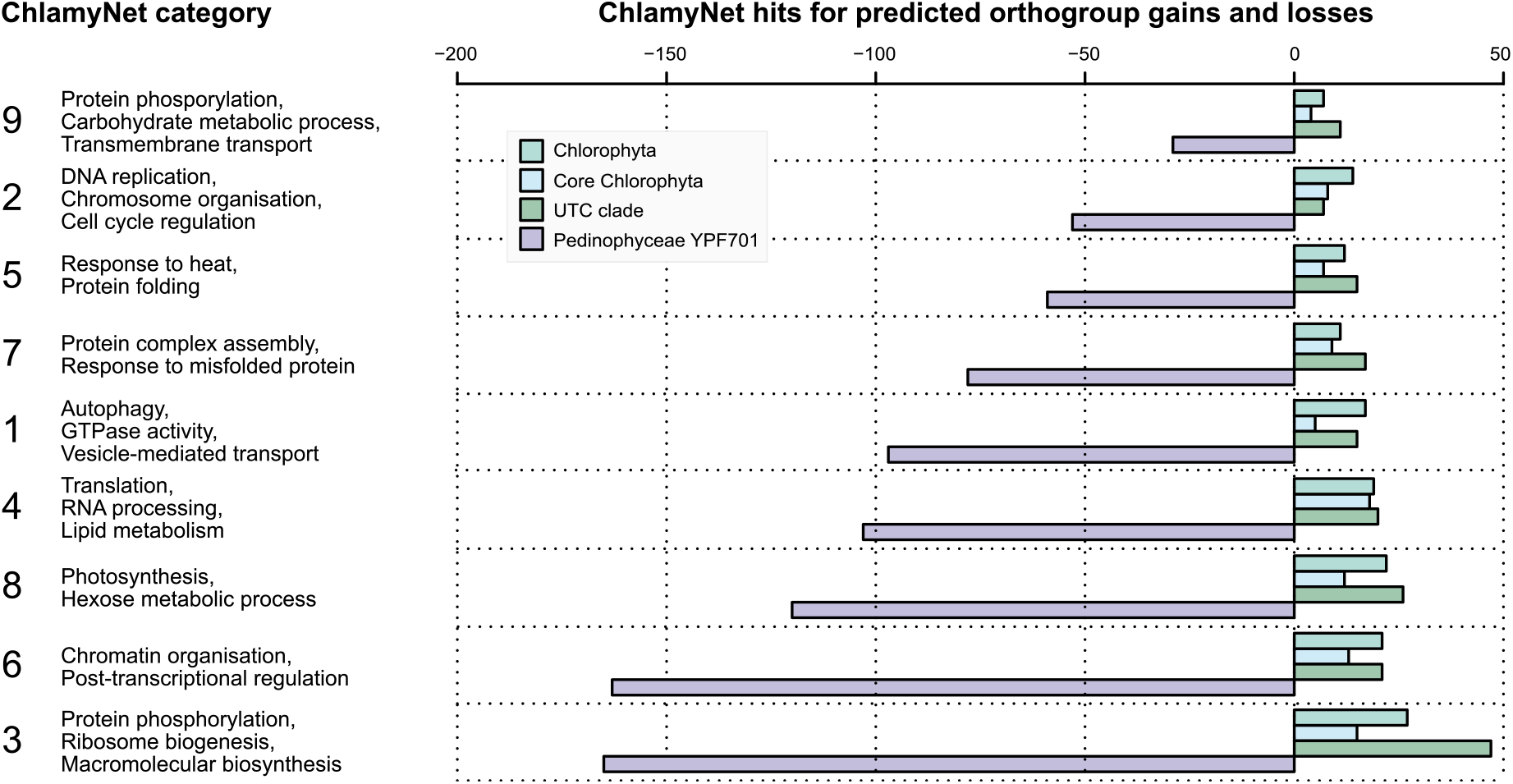
Number of orthogroups, containing *Chlamydomonas* sequences, gained along the branches leading to the Chlorophyta (Chloro), Core Chlorophyta (CC) and UTC clade (UTC), and lost along the branch leading to pedinophyte YPF701 (YPF), categorised into the 9 ChlamyNET gene clusters. y-axis labels refer to Gene Ontology (GO) term enrichment results for these clusters.

Considerable loss of orthogroups is predicted for the YPF branch following divergence from the rest of the Core Chlorophyta. A similar reduction of orthogroups is predicted for branches leading to each of the major prasinophyte groups, as well as to some individual taxa within the UTC clade (e.g. *Picochlorum, Ulva, Caulerpa*+*Ostreobium*). Sequences in orthogroups lost along the pedinophyte branch are diverse in function (Figs 3, S1), and many appear to be implicated in cell cycling and division, gene expression, and also include some light-related GO terms: “response to high light intensity”, “photosystem I assembly” and “red or far-red light signaling pathway” (Table S2). None of the orthogroups associated with these terms appear to be essential in the Chlorophyta, and they are absent in several lineages.

Conversely, a considerable gain in new orthogroups is predicted to have occurred along the UTC branch leading to the rest of the Core Chlorophyta following their divergence from the pedinophytes (Fig. 1). For the UTC, predicted gained orthogroups are associated with GO terms related to regulation, metabolism, reproduction and growth (Fig. 3, Table S2). In contrast, less change is predicted for the number of orthogroups along the branch leading to the last common ancestor of the Core Chlorophyta (CC) (Fig. 1, Table S3), and GO terms associated with predicted gained and lost orthogroups are reasonably balanced in terms of hierarchical clusters, lacking a clear functional pattern (Fig. S1, Table S2).

## Discussion

### Recurring genome minimisation in the green algae

The pedinophyte nuclear genome represents a missing link to examine early evolution of the Chlorophyta. The coding regions show strong codon and GC biases, which are also observed in their compact, intron-lacking chloroplast genomes (Marin, 2012; Jackson *et al*., 2018; Uthanumallian *et al*., 2021), indicating that they are under comparatively strong selection. The pedinophyte lineage also appears to have experienced considerable loss of homologous gene groups. These observations collectively support the hypothesis of selection for genome streamlining in the Pedinophyceae (Giovannoni, 2005). Genome streamlining appears to have occurred following the divergence of the Pedinophyceae from the rest of the Core Chlorophyta, with comparatively fewer changes in orthogroups predicted for the CC branch, and signals of genome reduction observed only for individual taxa within the UTC clade (eg. Gao *et al*., 2014; Foflonker *et al*., 2015). As Pedinophyceae are unicellular while many sequenced Core Chlorophyta are colonial and multicellular, the pedinophytes may have a larger effective population size, increasing the power of selection acting on their coding content to retain essential genes and remove non-essential DNA (Lynch, 2006; Smith, 2016).

Multiple independent signals of genome minimisation are observed at the base of the Chlorophyta: in the Pedinophyceae, Chloropicophyceae and Mamiellophyceae (Lemieux *et al*., 2019). This might indicate that the Chlorophyta common ancestor had a genome larger than those of many early-branching lineages. Evidence for larger ancestral Chlorophyta genomes remains circumstantial, but the conspicuous pattern of genome minimisation in early-branching lineages raises intriguing questions about the origin of Chlorophyta genomes. This predicted higher genomic novelty gained at deeper nodes followed by independent reduction events resembles patterns seen in recent comparisons of metazoan (Paps & Holland, 2018; Fernández & Gabaldón, 2020) and streptophyte (Bowles *et al*., 2020) genomes, and is consistent with the proposed universal biphasic model of speciation and genome evolution in eukaryotes, which involves initial rapid genome expansion (associated with emergence of new organism groups) followed by a prolonged period of gene loss driven largely by neutral processes and/or adaptive genome streamlining (Cuypers & Hogewe, 2012; Wolf & Koonin, 2013; Deutekom *et al*., 2019). The pedinophyte genome shows distinct features, high GC and stronger codon usage bias, which suggest coding content is under a different level of selection intensity compared with other reduced genomes found in the prasinophytes. It may be that prasinophyte lineages experienced lower relative selection intensity, thus different balances of natural selection and drift, during genome minimisation relative to the pedinophytes. Alternatively, prasinophyte lineages might have experienced a relaxation in selection following a period of streamlining; higher SCUO values for *Micromonas* relative to the rest of the Mamiellophyceae could be explained by lower relaxation of selection in this group, following the predicted genome minimisation in the Mamiellophyceae common ancestor (Worden *et al*., 2009). Differences between the genomes of early-diverging Chlorophyta lineages point to different balances of evolutionary forces driving their independent reduction events.

### Genomic innovation at the base of the UTC clade

Comparatively more orthogroup gains are predicted for UTC, following divergence of the pedinophytes, relative to the base of the Core Chlorophyta, suggesting a considerably high amount of genomic innovation arose along this branch. The highest average number of genes per homologous group for the Chlorophyta in our study were found in the UTC: for *C. lentillifera*, and fellow Ulvophyceae *Ostreobium quekettii* and *Ulva mutabilis* (Fig. 2), and members of the volvocine algae, whose relatively high gene duplication rates have been noted elsewhere (Hanschen *et al*., 2016). This reiterates the importance of gene duplication as a source of innovation in eukaryotic genomes (Wolf & Koonin, 2013). Bursts of conserved genomic novelty are attributed to whole-genome duplications (WGDs) in land plants (Bowles *et al*., 2020). However, studies investigating this phenomenon across the Viridiplantae have not identified evidence for WGDs in the ancestral branches of the UTC (One Thousand Plant Transcriptomes Initiative, 2019; Bowles *et al*., 2020).

Despite putative innovation in their common ancestor, considerable orthogroup loss is nonetheless predicted for many branches leading to individual taxa within the UTC clade. Although the nature of the Dollo parsimony method and false absences due to incomplete annotations might contribute to excessive losses being inferred (Wolf & Koonin, 2013; Deutekom *et al*., 2019), our results are consistent with the biphasic model of lineage genome evolution discussed above (Cuypers & Hogewe, 2012; Wolf & Koonin, 2013). The observed pattern of genome expansion in the common ancestor of the UTC followed by extensive gene loss within individual lineages may underpin the success of this species-rich clade in a diversity of habitats (Leliaert *et al*., 2012). Genomic innovation has the potential to open up many new niches for exploration by evolving organisms, while genome reduction is proposed to drive specialisation (Wolf & Koonin, 2013). Thus, new genes gained in their common ancestor may have provided genetic potential, which was then modified by lineage-specific patterns of evolution to enable the diversification of the UTC into many unique taxa inhabiting a diverse range of environments.

Our results from analysis of annotated gene functions reveal genomic novelty arising at the base of the Chlorophyta and UTC is implicated in a range of basic biological functions, from which specialised processes may have evolved. It appears that the genetic blueprint for many modern functions was already present in the Viridiplantae common ancestor. Probing these functional questions further is challenging, however, as the evolutionary emergence of many orthogroups predates the origin of the land plant-specific function based on their annotated GO terms (Leliaert *et al*. 2012; Romero-Campero *et al*., 2016; Bowles *et al*., 2020). A majority of orthogroups gained and lost along branches of evolutionary interest were not assigned GO terms (Table S3), meaning that the hypotheses proposed here represent merely the tip of the iceberg when it comes to study into the evolution of the functional gene repertoire of Chlorophyta.

Relative to work in the land plants and animals, comparative study into the evolution of Chlorophyta genomes is very much just beginning. This study represents merely an introductory peek into the diversification of green algae. Future study into Chlorophyta genome evolution would benefit from integration of genome annotations with functional work in order for inferences to be drawn about thus far uncharacterised genes. It is hoped that through initiatives striving for greater sampling within diverse Chlorophyta groups (e.g. Cheng *et al*., 2018), and parallel efforts to verify gene functions, the story of genome evolution in this lineage will continue to develop in the coming years.

## Acknowledgements

Joana Costa helped with DNA extraction for Nanopore sequencing. Nanopore sequencing was performed by Louise Judd. Ryan Wick generously aided with quality filtering of Nanopore reads and perspectives for their initial assembly. This work benefited from helpful comments by Geoffrey McFadden and Patrick Buerger on preliminary results included in S.I.R’s honours thesis. Funding for this work was provided by the Australian Research Council (DP150100705 to HV and CXC).

## Author Contribution

HV, CXC, CJJ and SIR designed the research; SIR, CI, CJJ, KU and YC performed the research; SIR, CI, KU and HV performed data analysis and interpretation; SIR, CI, KU, CJJ, YC, CXC and HV wrote the manuscript.

## Data Availability

The genome sequence of Pedinophyte YPF701 is available at the European Nucleotide Archive (ENA) with the project accession number ENA: PRJEB47395 and sample number ENA: ERS7299077. The raw Illumina reads are available with the accession numbers ENA: ERR6667563-ERR6667566, and the raw Nanopore reads are available with the accession number ENA: ERR6667567. Transcriptome reads are available with the accession number ENA: ERR6667568. The assembled genome is available with the accession number ENA: ERZ3455784.

## Supplementary Legends

**Fig. S1** CirGO visualisations of REVIGO results for GO terms associated with sequences in predicted gained orthogroups along branches Chloro, CC, UTC and YPF, and predicted lost orthogroups along branches CC, UTC and YPF, weighted according to the number of gained/lost orthogroups associated with each GO term.

**Table S1** Comparison of Viridiplantae nuclear genomes used for analysis in this study

**Table S2** Results of REVIGO analysis of GO terms associated with sequences in orthogroups predicted to be gained or lost along branches of interest.

**Table S3** Orthogroups predicted to have been gained and lost along branches of evolutionary interest that were the focus of this study, estimated using the Dollo parsimony principle, and the number of these orthogroups assigned GO terms by REVIGO, and containing a *C. reinhardtii* sequence associated with a ChlamyNET cluster.

