## Supplementary figures and images for "Nuclear genome of a pedinophyte pinpoints genomic innovation and streamlining in the green algae"

### Fig. S1

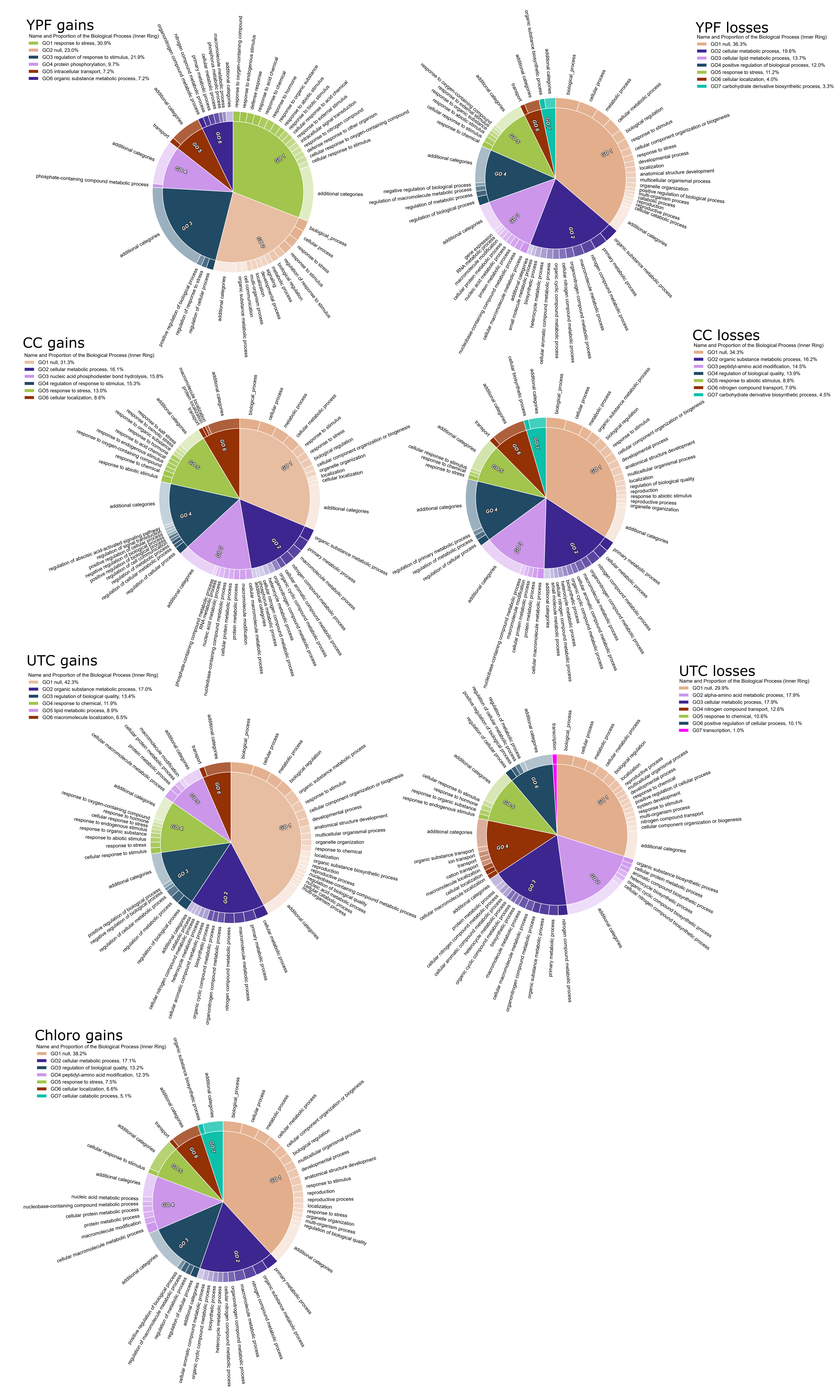
